# Neurophysiological correlates of collective perceptual decision-making

**DOI:** 10.1101/601674

**Authors:** Kristina G. Baumgart, Petr Byvshev, Alexa-Nicole Sliby, Andreas Strube, Peter König, Basil Wahn

## Abstract

Humans frequently coordinate with others in daily life. A recent study on perceptual decision-making showed that dyad members with similar individual performances attain a higher joint performance than the better dyad member (i.e., a collective benefit). However, little is known about the physiological basis of these results. Here, we replicate this earlier work and also investigate the neurophysiological correlates of decision-making using EEG.

In a two interval forced choice task, co-actors individually indicated presence of a target stimulus with a higher contrast and then indicated their confidence on a rating scale. Viewing the individual ratings, dyads made a joint decision. Replicating earlier work, we found a positive correlation between the similarity of individual performances and collective benefit.

We analyzed event related potentials (ERPs) in three phases (i.e., stimulus onset, response, and feedback) using explorative cluster mass permutation tests. At stimulus onset, ERPs were significantly linearly related to our manipulation of contrast differences, validating our manipulation of task difficulty. For individual and joint responses, we found a significant centro-parietal error-related positivity for correct versus incorrect responses, which suggests that accuracy is already evaluated at the response level. At feedback presentation, we found a significant late positive fronto-central potential elicited by incorrect joint responses, suggesting a stronger emotional response to negative as compared to positive feedback. In sum, these results demonstrate that response- and feedback-related components elicited by an error-monitoring system differentially integrate conflicting information exchanged during the joint decision-making process.

## 1. Introduction

In the words of Lynn Collins popularized by Rob Base and DJ EZ Rock, “It takes two to make a thing go right”. In a world inundated by such popular culture references, it is evident that there is a common socio-cultural belief in the benefits of collaboration. Accordingly, the dynamics of collaborative behavior are commonly studied to examine how groups working together have the potential to achieve higher performance than the best individual member in a group (i.e., collective benefit) (Bahrami et al., 2010, 2012a, 2012b; Fusaroli et al., 2012; Pescetelli et al., 2016). Research investigating group collaboration and its benefits encompasses many domains such as problem solving (Laughlin et al., 2002, 2006; Trouche et al., 2014), motor performance (Ganesh et al., 2014; Masumoto and Inui, 2013; Wahn et al. 2016a; Wahn et al., 2018b), and perceptual tasks (Bang et al., 2014; Mahmoodi et al., 2015, Wahn et al., 2018a,c). These studies suggest that information exchange and group dynamics are factors contributing to group benefit.

Perceptual decision-making paradigms are particularly suited for further investigation of factors affecting collective benefit, such as the type and quality of information exchanged and performance similarities (Bahrami et al., 2010, 2012a, 2012b; Brennan et al., 2008; Wahn et al., 2016b, 2017, 2018a, 2018c). Over a series of studies, Bahrami and colleagues discuss and examine the role of joint decision-making and collective benefit within a perceptual decision-making paradigm (Bahrami et al., 2010, 2012a, 2012b). Dyads were tasked with individually identifying a target stimulus from one of two intervals of circularly arranged Gabor patches. In conditions of dyadic disagreement, dyads either reached a consensus verbally or by means of confidence sharing when communicating nonverbally. Bahrami et al. (2010) found a collective benefit between individuals when allowed to speak freely about the task, and a trend towards this benefit (Bahrami et al., 2012b) when communicating nonverbally through a confidence rating scale.

Several models have been proposed to assess such collective behavior in perceptual decision-making tasks. Using a signal-detection model, Sorkin et al. (2001) investigated group dynamics and performance in relation to the individual and proposed an optimal model. In this model all information is available and is assumed to be perfectly exchanged between individuals, thus statistically setting the upper boundary of the collective benefit that can be achieved. Bahrami et al. (2010) argued that in a real-world setting, joint decision-making bears different constraints and limitations, whereby collective benefit can be reduced or disappear entirely under certain conditions. For example, when there are profound differences in sensitivity within a pair, suboptimal weighting of information occurs, resulting in a reduction of collective benefit (Bahrami et al., 2010). Hence, Bahrami et al. (2010) proposed the Weighted Confidence Sharing (WCS) model as a realistic alternative, accommodating for possible shortcomings in the exchange of information.

The physiological mechanisms of joint action and decision-making have also been investigated and insights have been used to establish a framework for understanding neurophysiological correlates of group tasks (Kourtis et al., 2013; Picton et al., 2012; Sebanz et al., 2006; Tsai et al., 2006). In regard to the physiological mechanisms of perceptual decision-making, there are a few key components which are particularly valuable to investigate. Attentional processes and visual awareness can be evaluated during stimulus presentation. The source of top-down biasing signals in visual perception derives from a network of areas in the frontal and parietal cortices (Kastner & Ungerleider, 2000). Parietal components are reported to be primarily concerned with the spatial component of perception, while frontal components are related to target detection, alerting, and motor representation (Kanwisher & Wojciulik, 2000). Components corresponding to visual awareness are also commonly investigated. These include the N200, a contralateral posterior negativity 200 ms after stimulus onset (Koivisto and Revonsuo, 2010), and the P300, a later positivity 300 ms after stimulus onset indicative of global attention processes leading to change detection (Eimer & Mazza, 2005). Furthermore, response and response conflict can be evaluated by the Readiness Potential (Bereitschaftspotential; Kornhuber & Deeke, 1965). Specifically, related to the collective decision-making task introduced earlier (Bahrami et al., 2010), for making the final joint decision it is assumed that the information of the partner is taken into account. Conflicting partner information could lead to a response conflict, which is associated with modulations of the readiness potential (Hackley & Valle-Inclán, 1999). In a review by Bartholow (2010), the readiness potential has been specifically discussed as being of interest in social-cognitive tasks involving conflict or the need for cognitive control.

Several other components have been proposed to be associated with response-error and performance monitoring directly after a response has been given. The term Error-Related Negativity (ERN) is used for a frontal negativity directly following response errors (Falkenstein et al., 1990). The ERN has been initially linked to error detection mechanisms, but ERN-like activity has also been found in correct trials (Bartholow et al., 2005). With this new evidence of correct-related negativity, the frontal response negativity stemming from the anterior cingulate cortex has instead been proposed as a response to fulfill a conflict monitoring function (Botvinick et al., 2001). Additionally, in response-error monitoring, an error positivity (Pe) occurring after the ERN has been consistently found (Overbeek et al., 2005). Some research suggests that the Pe may reflect the emotional significance of an error and is generally larger when participants are consciously aware that an error has been made (Orr and Carrasco, 2011). Moreover, Error-Related Feedback Negativity (fERN), which is similar to the response ERN and elicited after incorrect feedback (Miltner et al., 1997), is useful in evaluating error-monitoring.

Many researchers have investigated the neurophysiological correlates of joint action (Kourtis et al., 2013; Picton et al., 2012; Sebanz et al., 2006; Tsai et al., 2006). Many others (Bang et al., 2014; Fusaroli et al., 2012; Mahmoodi et al., 2015; Pescetelli et al., 2016) have extended the paradigm by Bahrami et al. (2010). However, none have yet quantified this particular behavior on a neurophysiological level. Therefore, the current study aims to bridge the gap by adapting the non-verbal condition of the Bahrami et al. (2012b) perceptual decision-making paradigm with the addition of EEG measurements. We first sought to adapt and reproduce the behavioral results of the paradigm. In particular, Bahrami et al. (2012b) found that as the sensitivity similarity between dyad members increases, the ratio of sensitivity between dyad and best member increases. As a second step, we assessed the metacognitive sensitivity of co-actors, which is a measure of the relationship between confidence and accuracy (Fleming and Dolan, 2012; Fleming and Lau, 2014; Pescetelli et al., 2016). Earlier studies found that there is a significant positive relationship between performance and confidence scale rating; the more confident the rating, the greater the performance. In the present study, we tested whether this is also the case for the present design. Bahrami et al. (2102b) compared results against model predictions of the proposed WCS and optimal models, finding neither model as a suitable predictor of joint performance under the parameters set for the study paradigm. In line with these analyses conducted by Bahrami et al. (2012b), and to further evaluate the possibility of achieving collective benefit, we compared the dyad performance with predictions of the WCS and optimal models.

Related to the neurophysiological correlates of collective decision-making, the paradigm by Bahrami et al. (2012b) offers a suitable design to investigate the neurophysiological underpinnings of collective perceptual decision-making. Specifically, the design of the paradigm allows for the comparison of individual and joint decisions revealing neurophysiological and behavioral insights of a joint decision from start to completion. Additionally, the nonverbal aspect of the paradigm allows for the use of EEG. Obviously, here some aspects of the original paradigm had to be adapted to accommodate the needs of the neurophysiological measurements. We exploratively examined electrophysiological potentials elicited by the fronto-parietal attention network and response- and error-related systems in the following phases: stimulus presentation, response, and feedback. First, we assessed attentional processes at the presentation of Gabor patches in the stimulus presentation phase. Secondly, we examined the Readiness Potential (RP) and the potentials related to response-error such as the ERN and the Pe at individual and joint responses in the response phase. Finally, we investigated potentials at the presentation of feedback as they may provide information about the emotional response to feedback regarding correctness of the response made by oneself or by one’s partner, and a conflict of these responses.

## 2. Methods

### 2.1. Participants

Participants were recruited from the student body of the University of Osnabrück, Germany. A total of 52 participants (33 female, age range 19-31 years) participated in the experiment after giving informed consent. All participants were healthy with normal or corrected-to-normal vision. Six pairs were excluded from analysis due to performing below chance level or failure to complete all experimental trials. Thus results are reported from 40 participants, comprised of 20 dyads, where 62.5% of participants reported previously knowing their partner. Each dyad consisted of one co-actor for whom we measured EEG (“EEG participant”) and one co-actor for whom only behavioral data was collected (“non-EEG participant”). Participants were recruited by departmental email and independently recruited for each role of the dyad. Participants could request to be paired with a friend and requests were accommodated when availability of both parties overlapped. Participants received either monetary compensation or course credit for their time. The study was approved by the ethics committee of the University of Osnabrück.

### 2.2. Experimental Setup

The current study adapted a nonverbal signal detection paradigm from Bahrami et al. (2012b), involving a Two-Interval Forced Choice (2IFC) design. Dyads were seated in a dark experimental room at the same table approximately 60 cm from individual display screens (BenQ 24", resolution = 1920 × 1080, 60 Hz refresh rate). Displays were controlled via PsychoPy 1.85.1 (Peirce, 2008) for Python 3.5 (Python Software Foundations). A divider was placed between participants such that they were unable to view their co-actor’s display. As both displays were connected to the same graphics card, an additional black cardboard cover was used to mask one half of the screen. For technical reasons, the EEG participant used the left half of the left display while the non-EEG participant used the right half of the right display.

Participants were each assigned a color to represent their corresponding tasks and responses; the EEG participant was blue and the non-EEG participant was yellow. Responses were made using the right hand on individual response boxes. Participants were provided with earplugs to reduce external noise interference and were instructed not to talk. The experimental phase contained 16 blocks of 16 trials after completion of one training block of 16 trials, which allowed participants to become familiar with the task. Experimenters remained present during training to ensure participants understood experimental instructions. Completion of both the training and experimental phases required approximately one and a half to two hours. A fifteen-minute break was given after the eighth block and additional breaks could be taken when necessary.

### 2.3. Stimuli

Stimuli were set to match those previously presented in the Bahrami et al. (2012b) study. The stimuli consisted of two sets of six vertically oriented Gabor patches (SD of Gaussian envelope = 0.38°, spatial frequency: 1.8 cycles/°, contrast: 10%), which were presented in circular formation (radius = 6°) equidistant from each other. Target stimuli were generated by increasing baseline contrast of a single Gabor patch in one of the following increments: 1.5%, 3.5%, 7%, and 15%. Location and interval in which the target stimulus appeared was randomized throughout the experiment. Contrast of the target stimulus was pseudorandomized such that all four contrasts occurred equally within an experimental session. Therefore, the difficulty of the task was randomized within a dyad such that all dyads experienced an equal number of trials per difficulty within an experimental session.

### 2.4. Procedure

A white fixation cross-marked the start of each trial. The non-EEG participant initiated each trial with a button press. The white fixation cross then turned black and was presented alone for a randomly jittered amount of time. After which, the first stimulus interval was presented for 85 ms, followed by a single black fixation cross for 1000 ms, and the second stimulus interval for 85 ms (Fig. 1). Participants then made a private individual “Interval Choice” indicating where the target stimulus occurred using a single button press, left for the first interval and right for the second.

**Figure 1.**
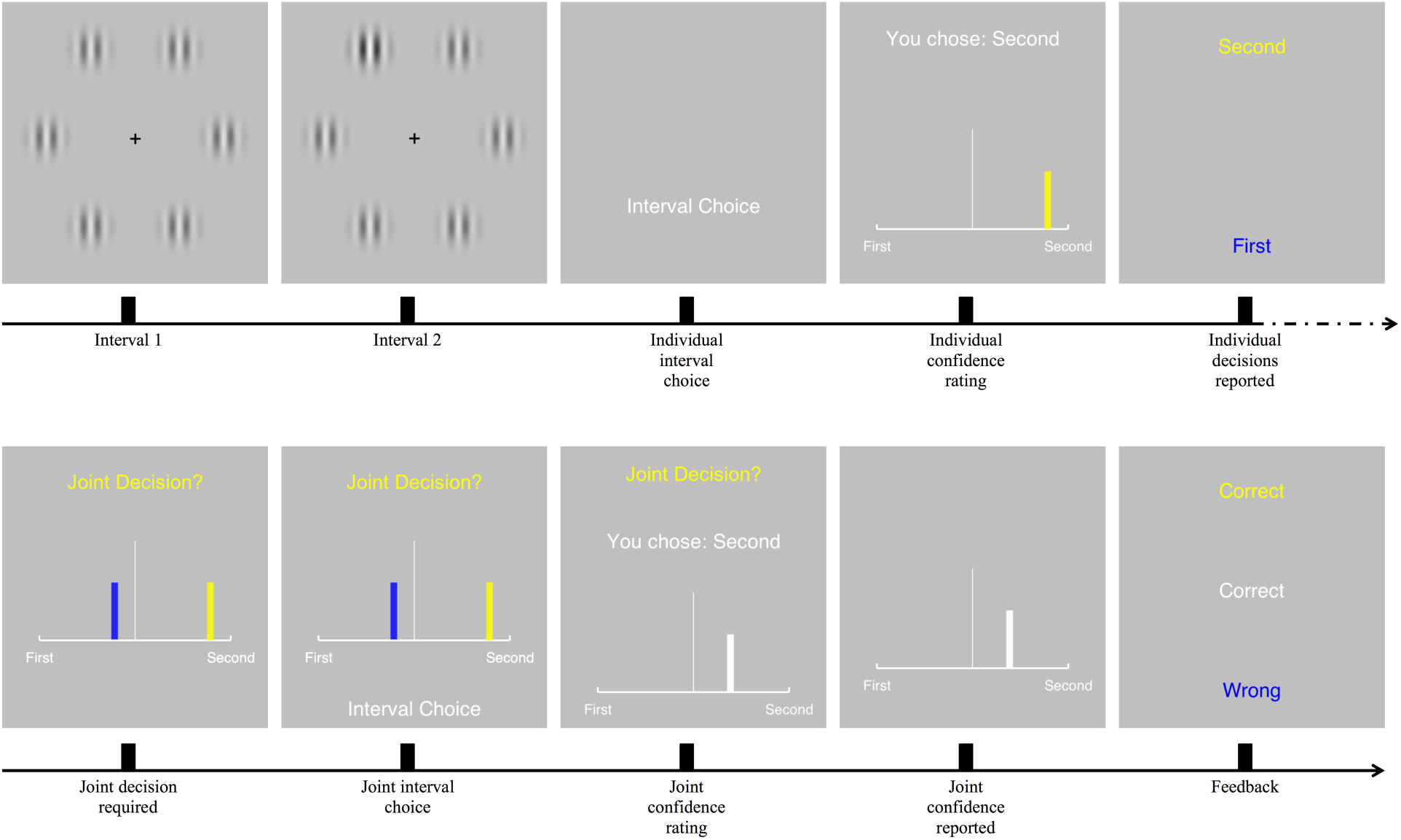
Procedural task and stimuli. Each trial began with the presentation of two stimulus intervals, one containing a target differing by contrast. A private button press indicating in which interval, first or second, the target appeared was made following stimulus presentation. Private confidence ratings were then made by moving the marker left for the first interval and right for the second interval with corresponding buttons on the response box. The farther from the central line, the more confident the decision. Individual interval choices were shared and a joint decision was made regardless of agreement or disagreement. One participant, indicated by instruction color, made a private joint interval choice and corresponding new private confidence rating. In the example above, the participant selected a slightly lower confidence rating for the joint decision as compared to her individual decision, presumably as the partner had chosen a conflicting decision at low confidence. Joint confidence rating was shared and feedback was given.

**Figure 2.**
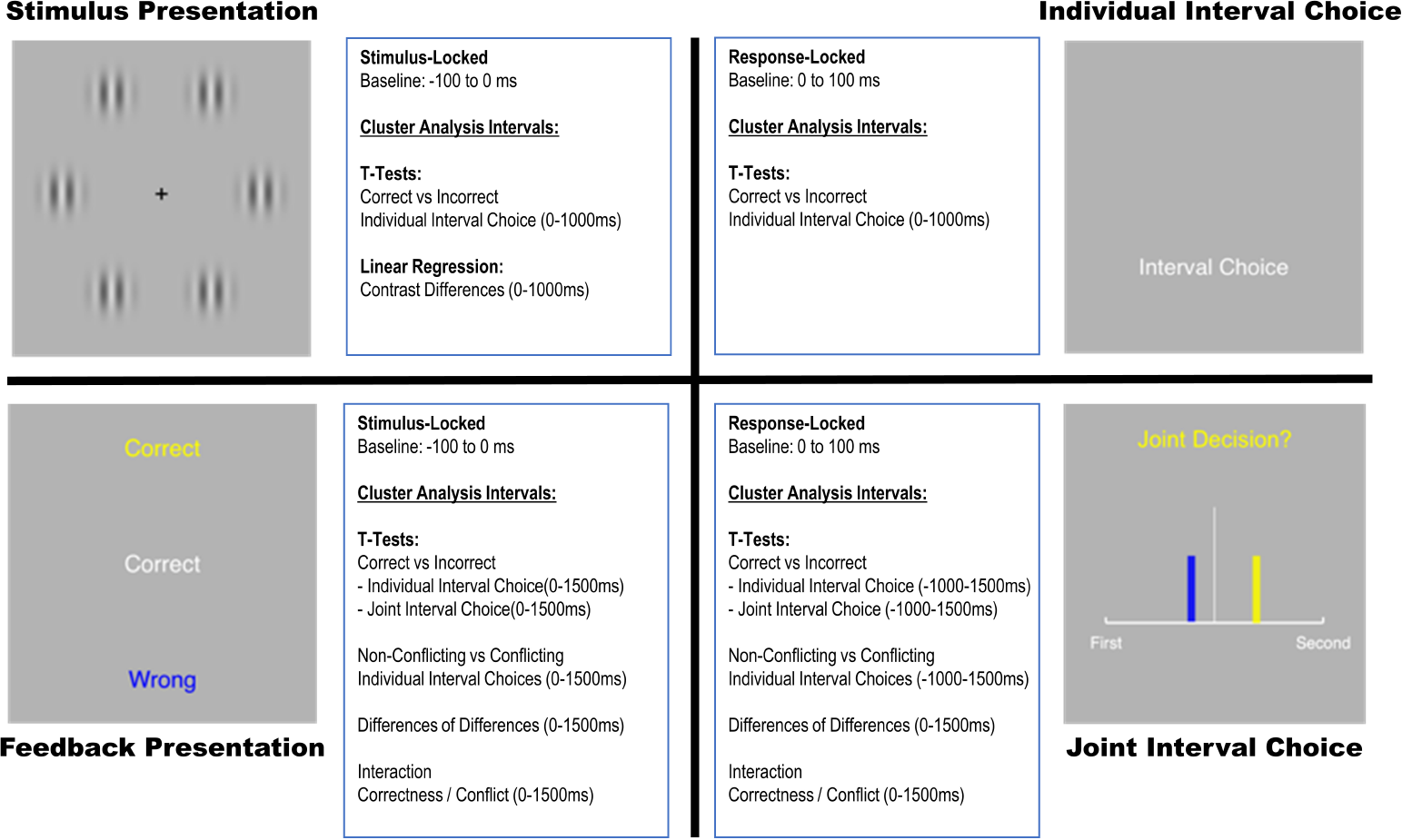
Information about EEG data processing (baseline and event lock) as well as statistical tests performed are listed for each respective phase. For each test, cluster analysis intervals are specified in brackets.

Following this button press, a bidirectional confidence rating scale was presented requiring participants to indicate confidence in their interval choice. A marker in the respective color of the participant was included in the center of the scale. Participants were forced to choose an interval as defined by the 2IFC design, and were only allowed to indicate confidence for their chosen interval. Private confidence ratings were made by moving the marker left for the “First” interval and right for the “Second” interval. The marker could be moved five increments in either direction; the farther from the central line, the more confident the decision.

After confidence rating submissions, individual interval choices were shared in the participants’ respective colors at the top or bottom of the screen. To avoid spatial bias (i.e., attention is allocated to a specific section of the display), the position of individual choices on the screen was randomized. Following this, individual confidence ratings were displayed and one dyad member, indicated by instruction color, was randomly chosen to make a joint decision for the dyad. As enacted in the individual decision, the joint decision consisted of an interval choice followed by a confidence rating. This confidence rating for the joint decision was chosen by the same participant who performed the joint decision. After the joint interval choice was made, the chosen interval was revealed to both participants. Both the interval choice and confidence rating were displayed after the respective choice. A black fixation cross was presented for 500 ms, followed by presentation of individual and joint performance feedback for 2000 ms. Joint feedback was always presented in white text in the center of the screen while individual feedback was presented in the participants’ respective colors in randomized positions (top or bottom). Feedback informed participants whether their joint and individual decisions were correct or not (i.e., “CORRECT” or “WRONG” was displayed on the screen). After feedback offset, a white fixation cross signaled the end of the trial and the start of a new one.

### 2.5. Behavioral Analysis

Behavioral analysis investigated factors contributing to collective benefit and compared the fit of WCS and optimal models, adapted as closely as possible from Bahrami et al. (2010, 2012b) and Sorkin et al. (2001). Contrast difference of the target stimulus, which varied by trial, was used as a measure of difficulty per trial to calculate a psychometric curve separately for joint and individual performance. The slope of the curve was then used to measure individual and dyad sensitivities. As in Bahrami et al. (2012b), performance was quantified as the proportion of second interval choices per reported locational contrast difference, which was positive if the target appeared in the second interval and negative if it appeared in the first. Collective benefit, which determined whether dyads achieved group benefit or not, was defined as the ratio of the dyad slope (*S*_*dyad*_) to the more sensitive participant (i.e. the participant with the higher slope, *S*_*max*_). Values above one indicate a collective benefit.

To further investigate factors contributing to collective benefit, participants’ use of the confidence rating scales were analyzed. We assessed in two ways how the similarity of confidence scale usage between co-actors of a dyad might contribute to collective benefit. The first approach compared the individual confidence value distributions of a dyad, defining a Distribution Difference Index (DDI), which quantifies the difference between the two distributions. This measure assessed scale-usage similarity as the difference in the number of times each respective confidence level was used between dyad members. The DDI was plotted against collective benefit on a per dyad basis. A DDI value of zero would indicate perfectly identical usage of all confidence levels. The DDI is expressed as

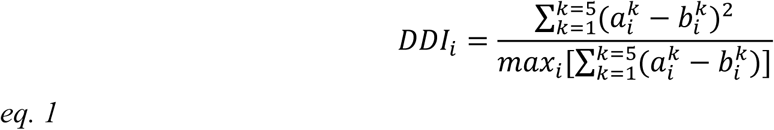

where 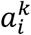 and 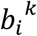 represent the proportion of answers given by each subject in the i-th pair at the k-th confidence level.

The second approach quantified how similarly dyads performed at any confidence level. To do this we proposed the Confidence Performance Difference Index (CPDI), which captures the difference in using confidence levels by the summed squared deviations. The individual performance of each participant was plotted against expressed confidence rating per trial. Then the difference vector for a dyad was calculated to evaluate the CPDI. A CPDI value of zero would indicate perfectly identical performance across all confidence levels. The CPDI, a variation of the DDI, is expressed as

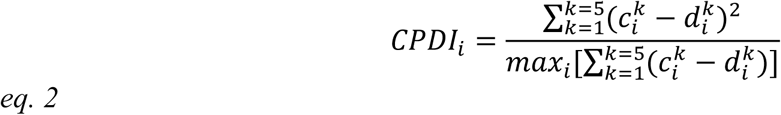

where the *CPDI*_*i*_ is a Confidence Performance Difference Index of i-th pair of subjects, 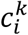 and 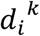 represent accuracy of subjects in the i-th pair at the k-th confidence level. Compared to more sophisticated measures like the Kullback Leibler Divergence it avoids the complexities of handling division by zero, as is the case with the Kullback Leibler Divergence when one category is not used by a participant at all. Furthermore, it has the advantage of simplicity making it transparent to non-experts as well.

Additionally, we sought to compare dyad performance in the present task to the performance predictions of the WCS and optimal models. The optimal model (Sorkin et al., 2001; Bahrami et al., 2012b), predicts dyad sensitivity as

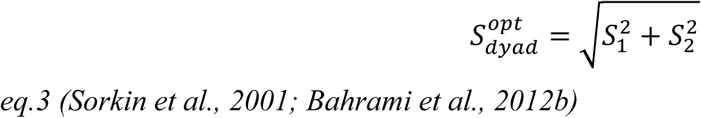

where *s*_1_ and *s_2_* represent the sensitivities of each individual. In this model, dyad sensitivity is expected to always be better than that of the best individual. To acknowledge differences in dyad information exchange, the WCS model is weighted by shared confidence, expressing dyad sensitivity as

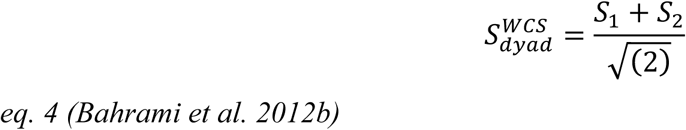

Given the equational variances, the models diverge further in instances of high dyad sensitivity difference.

### 2.6. EEG Data Acquisition

Data were recorded using asalab^TM^ with two 72 Refa amplifiers (ANT Neuro, Enschede, Netherlands) and gel-based Ag/AgCl 128 electrode cap (waveguard^TM^ caps), which were set based on the 10-5 international positioning (Oostenveld & Praamstra, 2001). For technical reasons, only 64 electrodes were used according to a 64 electrode cap layout. All recordings were referenced to the Cz electrode, with the ground positioned below the left clavicle utilizing a disposable cardiac electrode. Data were acquired at a 1024 Hz sampling rate and scalp impedances were kept below 10 kΩ. Eye movement was controlled through bipolar recordings of horizontal and vertical electrooculogram (EOG), made possible by placing disposable cardiac electrodes above and below the left eye, centrally aligned with the pupil.

### 2.7. EEG Preprocessing

Analysis and processing of EEG data were performed using the FieldTrip toolbox for MATLAB (Oostenveld et al., 2011). Mastoidal electrodes M1 and M2 were excluded from analysis due to excessive noise. The raw data were resampled to 256 Hz and Butterworth filtered at the third order at 0.1 Hz (high-pass) and 20 Hz (low-pass). The data set was epoched according to three phases, namely at the onset of Gabor patch presentation (stimulus presentation phase), at individual and joint responses of the EEG participant (response phase), and at the onset of feedback presentation (feedback phase). Epochs ranged from −2 to 2 seconds relative to the trial-defining trigger. These triggers were set at the onset of stimulus and feedback presentation as well as at the button press indicating individual and joint interval decisions. These epochs were then centered, i.e. the mean of each epoch was removed from the respective epoch. Outliers representing technical or movement artifacts were rejected manually after visual inspection. Subsequently, eye movement artifacts were corrected using independent component analysis (ICA) (Jung et al., 2000). Back-projection of the remaining non-artifactual components resulted in the corrected EEG data. All remaining trials with samples exceeding an absolute value of 80 µV in any EEG channel in a time frame from −1 to 1.5 seconds were deemed artifactual and were excluded from further analysis. For each subject and channel, the data was z-transformed, i.e. the mean of all samples of the respective subject and channel was subtracted from each sample and this difference was divided by the standard deviation. Please note that large inter-individual differences exist in EEG signals (Melnik et al., 2017). However, as a consequence of this transformation all subjects contribute equally to the grand average. The data were split into four different bins: 1) the onset of Gabor patches with a baseline from −100 to 0 ms from stimulus onset, 2) individual decisions with a baseline from 0 to 100 ms from the button press indicating the individual decision, 3) joint decisions with a baseline from 0 to 100 ms from the button press, and 4) the onset of feedback with a baseline ranging from −100 to 0 ms where all combinations of feedback were considered for analysis (Fi

### 2.8. EEG Statistical Analysis

Grand averages (GAs) were calculated by the unweighted mean of z-transformed data of each participant for each trial separately for each phase and feature of interest. For each phase, we define conditions that we aim to compare. Please note that condition is used only to separate experimental trials on the basis of certain criteria, not that these are literally under experimental control. In particular, at stimulus presentation and at the individual decision, a feature of interest is represented in the correctness of the response, i.e. conditions that will be compared were defined as trials with a correct or incorrect joint response. At feedback presentation and the joint decision button press, features of interests were the correctness of the individual and joint response and whether responses of participants conflicted. Hence, conditions are correct or incorrect individual and joint responses, respectively, and conflicting or non-conflicting responses. All statistical tests were conducted in electrode space and were corrected for multiple comparisons using non-parametrical permutation tests of clusters of neuronal activity (Maris & Oostenveld, 2007). Please note that this procedure allows a data driven investigation of the physiological substrate of joint decision processes without a bias to pre-selected electrodes. Importantly, in contrast to other exploratory procedures it handles multiple comparison corrections intrinsically and efficiently. For the present purpose it combines two important advantages that it is not dependent on a specific theoretical framework, but allows an unbiased investigation of the physiological decision process. Clusters were formed based on neighboring time-channel bins exceeding a parametric statistical test value corresponding to a p-value of 0.05. Clustering was restricted to samples having at least two neighboring channels also exceeding the threshold, where neighbors of channels were defined by a triangulation algorithm implemented in the FieldTrip toolbox. Using Monte Carlo resampling, 5000 matrices with randomly permuted condition labels within each subject were generated, as were corresponding clusters for each of these matrices.

For each cluster, cluster-level statistics were calculated using the cluster mass (Bullmore et al., 1999), which is the sum of statistical values of all samples included in a cluster. Clusters obtained from the original data matrix were compared with the permutation distribution of the largest cluster masses resulting from each random permutation of data labels. A p-value represents the proportion of clusters in random permutation exceeding the cluster mass value of those obtained from the original data matrix. This test was applied two-sided for negative and positive clusters separately and resulting p-values were Bonferroni corrected by the factor 2 for the two-sided test accordingly.

Using this procedure in the stimulus presentation phase, a linear trend of the four levels of contrasts was evaluated for data locked to the onset of the presentation of Gabor patches using a dependent samples linear regression test implemented in the FieldTrip toolbox for MATLAB (Oostenveld et al., 2011). Prior to this test, conditions were averaged according to the four levels of contrasts. The test was applied to samples in a time frame ranging from 0 to 1000 ms. Additionally, dependent sample t-tests were applied to compare conditions in different time frames. In particular, at the presentation of Gabor patches, conditions were averaged separately according to correct and incorrect individual responses for each participant and the cluster permutation test was applied in a time frame from 0 to 1000 ms.

In the response phase, correct and incorrect responses with EEG data locked to the button press indicating the individual decision were tested in a time frame from −1000 to 1500 ms. Data locked to the button press of the EEG participant indicating a joint decision were tested in four different ways. First, conditions were averaged according to conflicting and non-conflicting individual responses, where in conflicting trials individual responses differed while in non-conflicting trials they were the same. Secondly, conditions were averaged according to correct and incorrect joint decisions. Correct versus incorrect and conflicting versus non-conflicting contrasts were tested in a time frame ranging from −1000 to 1500 ms. Thirdly, the absolute difference of correct and incorrect responses versus the absolute difference of conflicting versus non-conflicting responses was assessed in a time frame ranging from 0 to 1500ms. Finally, an interaction of conflict and correctness was assessed using the same procedure, i.e. the absolute difference between conflicting correct and incorrect trials was tested versus the absolute difference between non-conflicting correct and incorrect trials in a time frame ranging from 0 to 1500ms.

In the feedback phase, conditions were averaged either according to feedback representing the correctness of individual response or to feedback representing the correctness of joint response. Correct and incorrect feedback conditions were tested for significant differences for individual and joint feedback separately. Furthermore, individual conflicting feedback was tested against individual non-conflicting feedback, where individual responses disagree or agree, respectively. These tests of feedback were done using the cluster-permutation procedure described above. Additionally, an interaction of individually conflicting conditions and correctness of the joint interval choice was tested using the same cluster-permutation procedure and was further quantified using paired-samples t-tests. All cluster permutation tests at feedback onset were applied in a time frame ranging from 0 to 1500 ms. ERP plots were obtained by averaging trials of a certain condition at first within a participant. Then, a Grand Average (GA) was produced by the unweighted average of the obtained averaged ERPs of each participant. These Grand Averages were plotted for visualization.

## 3. Results

### 3.1. Behavioral Results

Although this work focuses on the neurophysiological mechanisms supporting joint decisions, the experiments speak to questions on the behavioral level as well. Therefore, in the following section we report the behavioral results in close analogy to previous studies.

#### 3.1.1. Comparison of dyad with the better participant

Dyad sensitivity was compared to the sensitivity of the better participant to assess whether a dyad achieved collective benefit. On average dyad sensitivity was not significantly different from the sensitivity of the better participant (Fig. 3B left bar, effect size -0.1458, *t*(19) = -1.517, *p* = .146, one sample t-test). That is, pairs did not significantly attain a collective benefit. In spite of this, a significant relationship between the ratio of individual sensitivity similarity (*S*_*min*_/*S*_*max*_) and the ratio of sensitivity between dyad and best member (*S*_*dyad*_/*S*_*max*_) was found. As the ratio of similarity increases, the ratio between dyad and best member increases; therefore the possibility of achieving collective benefit also increases (Pearson *r* = 0.877, *p* < .001, *n* = 20).

**Figure 3.**
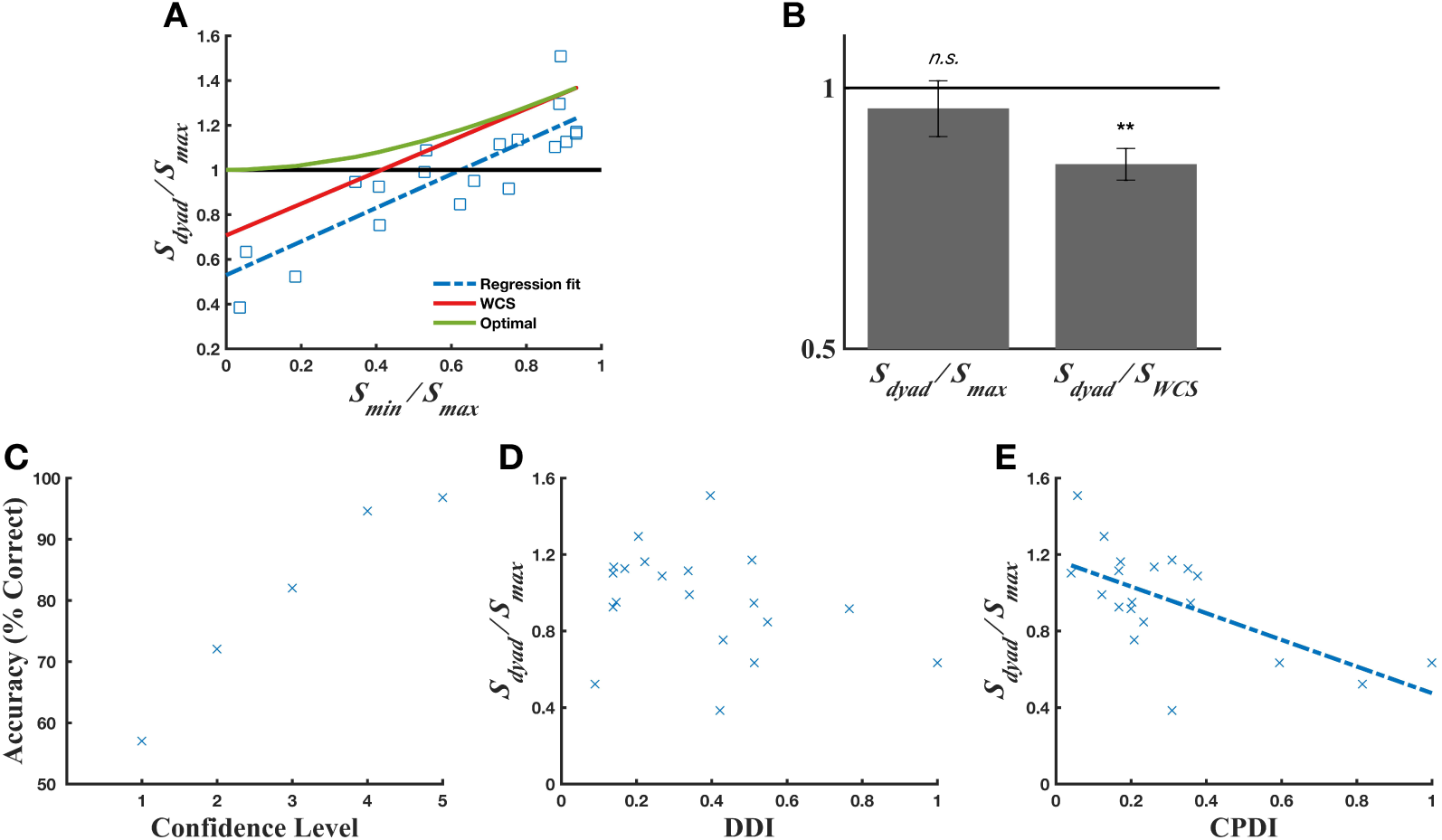
Behavioral results overview. **(A)** Model prediction comparison with experimental data. Dyad sensitivity (*S*_*dyad*_/*S*_*max*_) in relation to the ratio of individual sensitivities (*S*_*min*_/*S*_*max*_) is plotted for 20 pairs (blue squares) against the Weighted Confidence Sharing Model (WCS, red line) and Optimal Model (green line) predictions. A dyad sensitivity value above one indicates achievement of collective benefit to the point of joint advantage. **(B)** Optimality of group performance combined for all pairs compared against the better participant (left bar) and WCS model (right bar). Double asterisks indicate a p-value < .001. Error bars signify SEM (*n* = 20). **(C)** Average performance in relation to expressed confidence level for all participants. Each data point indicates performance accuracy for the stated level of confidence. One indicates low and five indicates high confidence. **(D)** The relationship between Distribution Difference Index (DDI) and collective benefit (*S*_*dyad*_/*S*_*max*_) per dyad. **(E)** The relationship between Confidence Performance Difference Index (CPDI) and collective benefit (*S*_*dyad*_/*S*_*max*_) per dyad (regression fit, blue dashed line).

#### 3.1.2. Confidence rating scale usage and performance vs. collective benefit

The relationship between performance and expressed confidence level was used as a measure indicating metacognitive sensitivity on an individual level (Fleming and Dolan, 2012; Fleming and Lau, 2014; Pescetelli et al., 2016). A positive relationship was found between performance and expressed level of confidence, where accuracy increased as expressed confidence level increased, thus indicating higher metacognitive sensitivity (Fig. 3C). Following this, the Distribution Difference Index (DDI) was used to assess how confidence scale usage might affect collective benefit. The DDI quantified this distribution by taking the difference in the number of times each confidence level between a dyad was used. The data do not suggest a significant relationship between the DDI and collective benefit (Fig. 3D, Pearson *r* = -0.3, *p* = *.19*, *n* = 20). That is, how differently a pair distributed their confidences across the rating scale did not predict collective benefit. Further investigation of confidence scale usage was assessed by the Confidence Performance Difference Index (CPDI). The CPDI quantified the difference in performance between a dyad for each expressed confidence level. Confidence level performance accuracy was used to calculate CPDI per dyad. The observed relationship between CPDI and collective benefit suggests that CPDI is a predictive measure of collective benefit in that, as CPDI increases the likelihood of achieving collective benefit decreases significantly (Fig. 3E, Pearson *r* = -0.62, *p* = .003, *n* = 20). Other measures, such as the KL-divergence, have been commonly used to assess differences in distributions, similarly to our DDI and CPDI measures. We therefore used the KL-divergence to test whether we would get similar results as for the DDI and CPDI. Indeed, we found that the KL-divergence also predicted the collective benefit similarly to the CPDI (Fig. S1, Pearson r = -0.64, p = .003, n = 20). Thus supporting similar previous findings, in order to achieve a collective benefit it is not decisive to use different confidence levels with similar frequency, but to use different confidence levels to indicate comparable levels of individual performance.

#### 3.1.3. Comparison of dyad with Weighted Confidence Sharing and optimal models

Dyad performance was compared to the predictions of the WCS and optimal models to further investigate the possibility of achieving collective benefit. The WCS model predicted significantly higher dyad performance than was observed (Fig. 3A-3B right bar, effect size 0.2456, *t*(19) = -4.82, *p* < .001, one sample t-test), although it seems to predict a similar linear dependency on performance similarity. Considering the optimal model predicts higher overall collective benefit in terms of relative sensitivity difference compared to the WCS, the optimal model also predicted significantly higher dyad performance than was observed (Fig. 3A, *t*(19) = - 5.3, *p* < .001, one sample t-test). Thus, in spite of the significant deviations, of the models tested the WCS model performs best and specifically describes a similar linear dependency on performance similarity.

#### 3.1.4. Correctness vs. Familiarity

Dyad familiarity was evaluated to exclude it as a confounding variable affecting achievement of collective benefit. In a post-experiment survey, participants were asked to what degree, if at all, they had previously known their partner. No significant difference in dyad performance was found based on reported familiarity (Pearson *r* = -0.12, *p* = .61, *n* = 20).

### 3.2. EEG Results

#### 3.2.1. Stimulus Presentation

We first sought to investigate cortical mechanisms at stimulus presentation. At the presentation of Gabor patches, no significant cluster representing a significant difference in EEG signals between correct and incorrect individual decisions was found (all *p*-values > .085). The linear regression statistics for the cluster analysis of dependent samples of different contrast levels at the presentation of Gabor patches revealed two significant clusters, one positive and one negative, corresponding to a dipole. The positive cluster included samples from electrodes in fronto-central to parieto-central positions, 430-871 ms after stimulus onset (Cluster Mass (CM) = 4530; *p* = .002; Confidence Interval of p (CI) = [0.0008; 0.0032]). The negative cluster included samples from left temporal areas 297-859 ms from stimulus onset (CM = -3975; *p <* .001; CI = [0.0002; 0.0016]). The highest parametric positive statistical t-value was found at channel AF4 at 488 ms, (Fig. 4 top, *t*(19) = 5.88, *p* < .001, *d =* 1.31). The highest parametric negative statistical t-value was found at channel FT7 at 535 ms (Fig. 4 bottom, *t*(19) = -6.75, *p* < .001, *d* = -1.51). These results indicate a modulation of the EEG signals by the experimental manipulation of contrast differences at a relatively late latency of topographically widespread event-related potentials.

**Figure 4.**
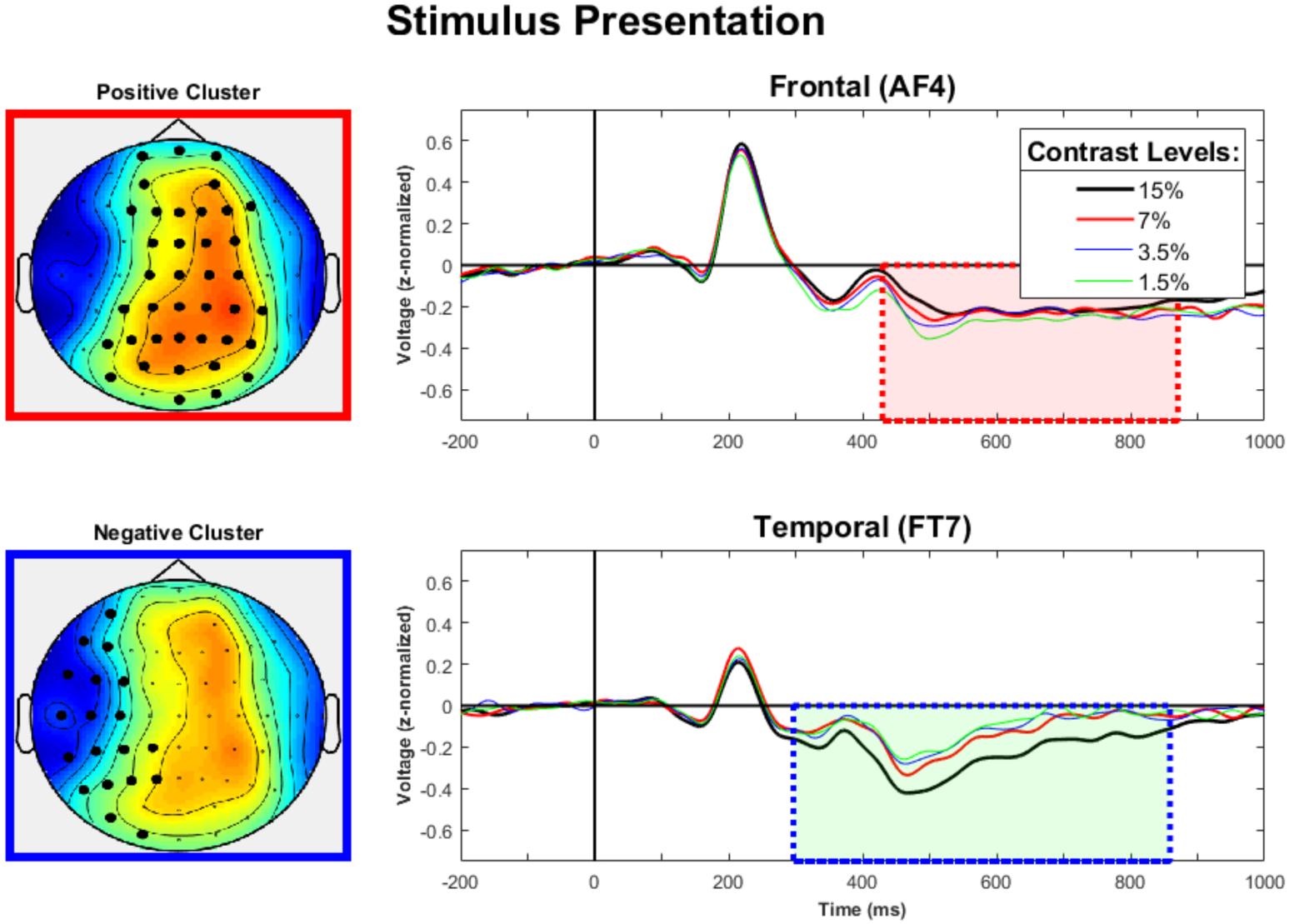
The averaged ERPs over each participant are shown for AF4 (top), and FT7 (bottom). Red and blue shaded boxes show significant time frames of the significant positive and negative clusters representing a linear regression of contrast differences, respectively. On the left, topographies of these clusters with averaged t-values are accordingly color-coded and significant electrodes are highlighted. Warm colors (yellow-red) represent positive t-values, cool colors (blue) negative t-values.

#### 3.2.2. Individual Decision

At the button press indicating the individual interval decision, we investigated whether EEG signals were modulated by the correctness of the response. The permutation test of clusters composed of t-values obtained from dependent samples t-tests revealed one significant negative cluster. Thus, indicating lower values for correct individual decisions, as compared to incorrect individual decisions, at central to fronto-central right hemispheric electrodes from 578-1097 ms after response (CM = -3296,6; p = .009; CI = [0.0062; 0.0114]). The highest parametric negative statistical t-value was found at channel C4 at 1018 ms, (Fig. 5, *t*(19) = -5.01, p < .001, *d* = -1.23). We can conclude that event-related potentials elicited after the individual response were modulated by the correctness of the response. No differences have been found before the response despite a time frame ranging well before the response trigger was tested, indicating that response preparation was not affected by factors of correctness or conflict.

**Figure 5.**
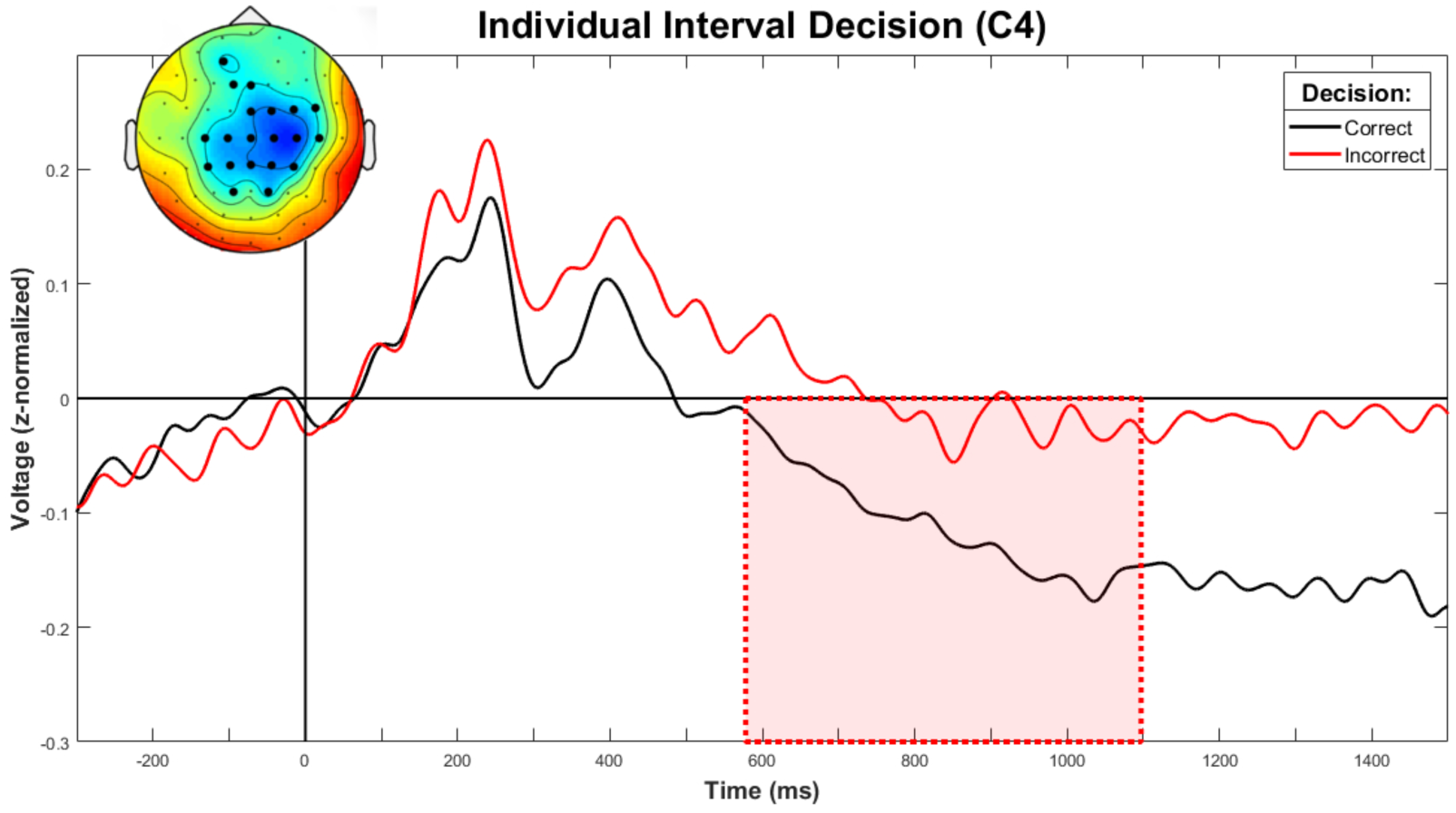
The averaged ERPs of correct and incorrect responses over each participant at the button press indicating individual interval choice are shown. The red shaded area represents the time frame of the cluster representing a significant difference between correct and incorrect trials. For this cluster, a topography representing averaged t-values over the significant time frame with highlighted electrodes is shown. Warm colors (yellow-red) represent positive t-values, cool colors (blue) negative t-values.

#### 3.2.3. Joint Decision

At the joint decision, we conducted tests to find manifestations of conflicting individual decisions as well as of the correctness of the joint decision within event-related potentials. Similarly to the reported difference after the individual decision, a significant negative cluster was found representing a significant difference between correct and incorrect joint interval decisions after the response of the EEG participant. This suggests lower values for correct joint interval decisions as compared to incorrect joint interval decisions at centro-parietal electrodes, 699-1223 ms after response (CM = -3426; p = .0156; CI = [0.0112; 0.019]). The highest parametric negative statistical t-value was found at channel CPz at 781 ms (Fig. 6, *t*(19) = -6.57, *p* < .001, *d* = -1.47). The permutation test of clusters representing differences between conflicting and non-conflicting conditions revealed two significant clusters, one positive and one negative. The positive cluster, with a broad frontal topography, indicated a higher value for non-conflicting as compared to conflicting conditions, 106-391 ms after response (CM = 3561.7; *p* = .0488; CI = [0.0428; 0.0548]). The highest parametric positive statistical value was found at channel AF7 at 195 ms (*t*(19) = 6.438, *p* < .001, *d* = 1.44). The negative cluster, corresponding to a higher value for conflicting conditions, was found at central to parietal electrodes, 82-1500 ms after response (CM = -21201; p < .001; CI = [0; 0.001]). The highest negative parametric statistical value was found at channel CP1 at 203 ms (*t*(19) = -7.54, *p* < .001, *d* = -1.69). Thus, both main effects of conflict, at frontal and parietal electrode sites, and correctness at parietal electrode sites, were found in the statistical test of event-related potentials. Again, only a time frame after the button press was found to be significantly affected by tested factors. We then tested whether those conditions differed in their magnitudes. In order to do so, the absolute differences of these conditions were tested for significant differences using the previously introduced cluster-permutation procedure.

**Figure 6.**
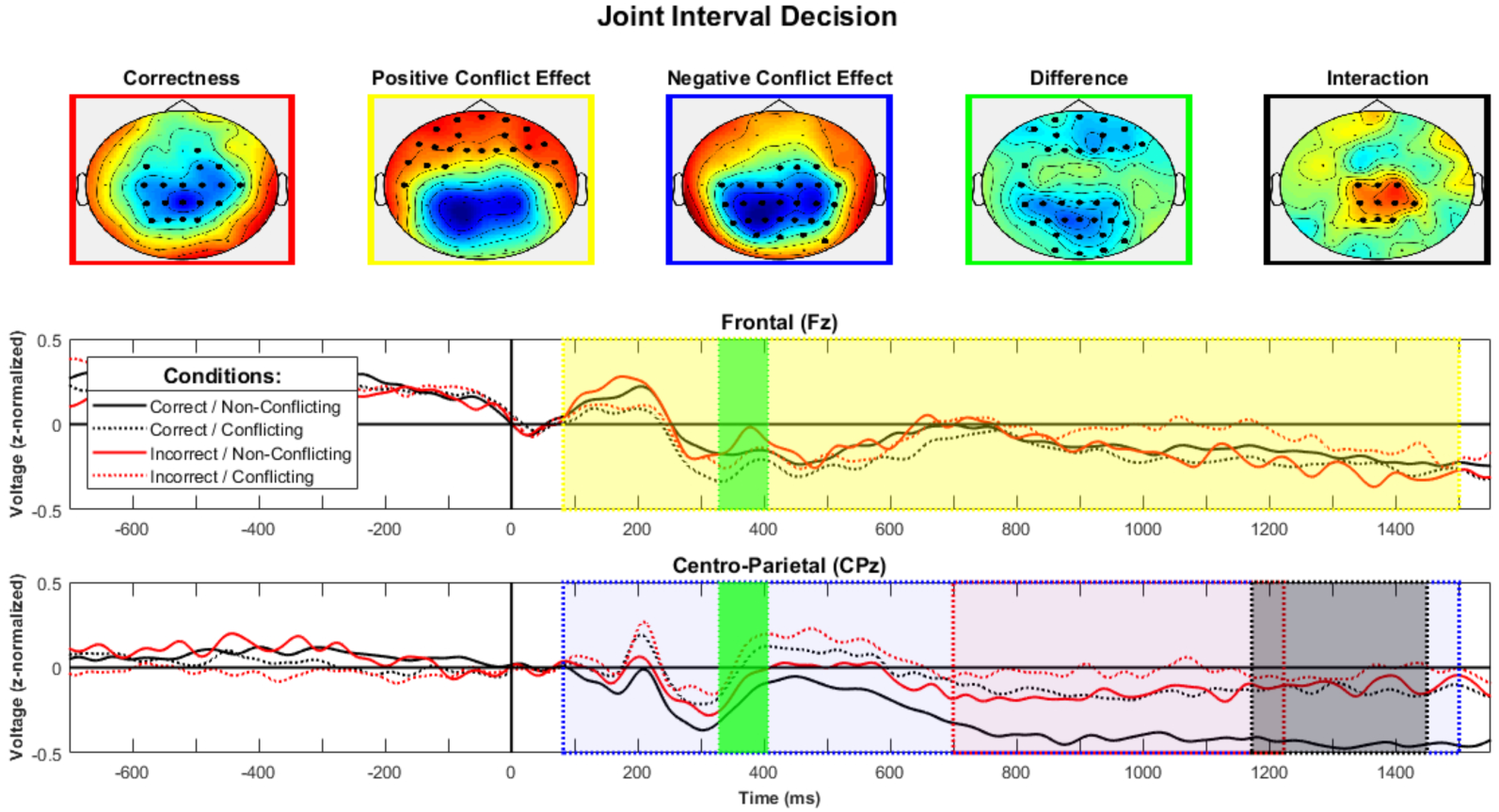
Average ERPs corresponding to conditions with different conflicting and different correctness situations of the joint interval choice are shown for Fz (middle) and CPz (bottom). Colored shaded areas show significant time frames of corresponding significant clusters. Red corresponds to the negative cluster representing differences between correct and incorrect trials. Yellow and blue correspond to the significant positive and negative clusters of conflicting situations. Green refers to the cluster testing the difference of the correctness contrast versus the conflict contrast and black represents the respective interaction. Topographies (top) are accordingly color-coded and represent averaged t-values over the cluster. Warm colors (yellow-red) represent positive t-values, cool colors (blue) negative t-values.

This test yielded one negative cluster representing a significantly lower absolute difference between correct versus incorrect as compared to conflicting versus non-conflicting joint interval decisions. Thus indicating a higher absolute difference between values of conflicting versus non-conflicting conditions as compared to correct versus incorrect conditions. This cluster included electrodes in frontal and parietal positions connected by a narrow bridging electrode in a time frame 328-406 ms after response (Fig. 6, CM = -771.7, *p* = .013; CI = [0.01; 0.016]). For this effect, the highest negative parametric statistical value was found at channel F2 at 340 ms (*t*(19) = -4.57, *p* < .001, *d* = -1.02). This test indicated that the conflict condition elicited a higher difference in event-related potentials at both frontal and parietal electrode sites.

Finally, we investigated whether the different correctness and conflict trials interacted. There was a significant interaction 1172-1445 ms after response, indicating higher differences between correct and incorrect interval decisions in non-conflicting conditions as compared to conflicting conditions (CM = 946.5, *p* = .005, CI = [0.003; 0.007]). This interaction was found at centro-parietal electrode sites. The highest parametric statistical value was found at channel C2 at 1293ms (Fig. 6, *t*(19) = 4.93, *p* < .001, *d* = 0.66). In conclusion, our results indicate differential representations of correctness and conflict in frontal and parietal event-related slow evoked potentials after the button press indicating the joint decision.

#### 3.2.4. Feedback

We evaluated three different conditions represented in the feedback display: (1) correctness of the individual decision, (2) correctness of the joint decision and (3) conflict of individual decisions. (1) No significant cluster was found representing differences between a correct and incorrect individual interval decision (all *p*-values >.07). (2) Furthermore, no significant cluster representing differences between feedback about conflicting and non-conflicting joint decisions was found (all *p*-values > .143). (3) A significant negative cluster was found for differences between feedback conditions, signaling a correct versus incorrect joint decision in trials where the EEG participant made the joint decision (CM = -5060; *p* = .004; CI = [0.002; 0.006]). This frontal to parietal cluster included samples in a time frame 449-883 ms after feedback onset. The highest negative parametric statistical value was found at channel FC1 at 645 ms (Fig. 7, *t*(19) = -6.52, *p* < .001, *d* = -1.46). This series of tests indicated that only the joint correctness condition elicited significant differences in event-related potentials after feedback presentation.

**Figure 7.**
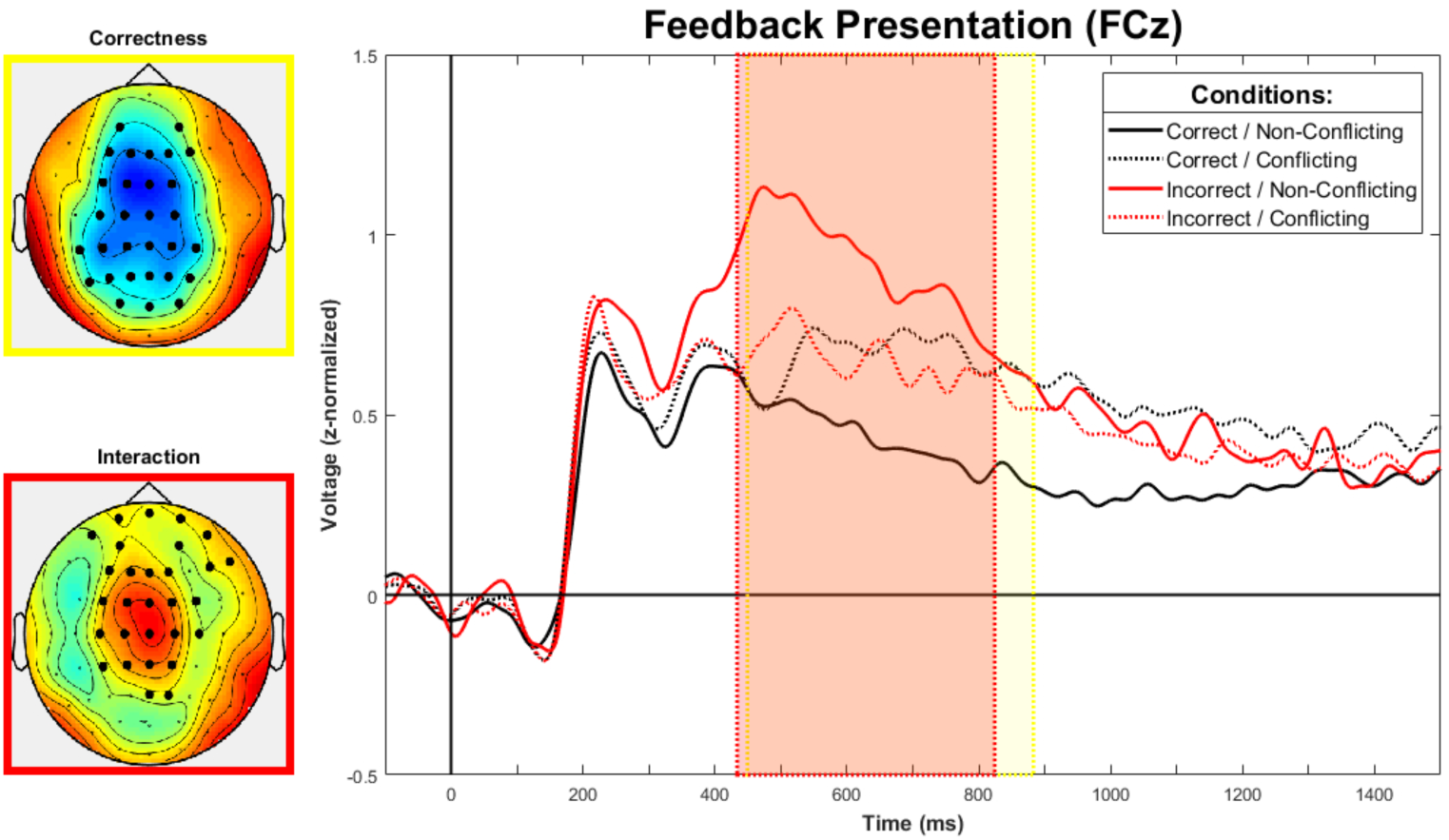
Average ERPs corresponding to conditions with different conflicting situations and the correctness of the joint interval choice are shown for FCz. The yellow shaded area represents the time frame of the significant cluster of the correctness contrast. The red shaded area represents the significant interaction. Topographies on the left are accordingly color-coded and represent positive (yellow/red) and negative (blue) t-values. Significant electrodes are highlighted.

Subsequently, we further investigated if this difference was modified by a conflict. We found a significant interaction 434-824 ms after feedback onset at frontal to central electrode sites, indicating higher differences between correct and incorrect joint interval decisions in non-conflicting conditions as compared to conflicting conditions (CM = 2799.2; *p* < .001; CI = [0; 0.0001]). The highest parametric statistical value was found at channel FCz at 793 ms (Fig. 7, *t*(19) = 6.55, *p* < .001, *d* = 1.46). In the same time frame, 434-824ms, both conflicting correct responses (*M* = 0.67, *SD* = 0.58; *t*(19) = 2.293, *p* = .0167 *d* = 0.71,one-sided) and conflicting incorrect responses (*M* = 0.66, *SD* = 0.53; *t*(19) = 2.73, *p* = .007, *d* = 0.73, one-sided) elicited significantly larger ERPs than non-conflicting correct responses (*M* = 0.48, *SD* = 0.33) at FCz. Additionally, conflicting correct responses (*t*(19) = -2.68, *p* = .007, *d* = -0.21, one-sided) and conflicting incorrect responses (*t*(19) = -3.64, *p* < .001, *d* = -0.26, one-sided) elicited significantly smaller ERPs than non-conflicting incorrect responses (*M* = 0.933, SD = 0.57) at the same channel and time. These results indicate that conflicting correct and conflicting incorrect responses evoked a significantly lower potential than non-conflicting incorrect decisions as well as a significantly higher potential than non-conflicting correct decisions. Finally, we investigated whether differences in event-related potentials, resulting from the initially tested conditions, differed from each other. The permutation test of clusters representing significant differences between differences of joint versus individual correctness, joint correctness versus conflict conditions, and of individual correctness versus conflict conditions yielded no significance (all *p* values > .213). In summary, these results show that feedback-related differences in EEG signals were elicited by incorrect versus correct responses.

## 4. Discussion

In the present study, we investigated factors contributing to and neurophysiological correlates of collective decision-making and collective benefit by adapting a perceptual decision-making task. Similarly to the nonverbal condition in Bahrami et al. (2012b), we found a trend towards collective benefit for dyads communicating via confidence rating scales. Furthermore, performance improved as expressed confidence level increased, indicating higher metacognitive sensitivity. Evaluation of confidence rating scale usage revealed no significant relationship between the DDI, the difference in the frequency with which each confidence level was used between a dyad, and collective benefit. However, the CPDI, the difference in performance at each confidence level between a dyad, was found to be a predictive measure of collective benefit. Neither the optimal nor the WCS model predictions significantly fit the data, yet the WCS model produced a similar linear tendency on performance similarity. Dyad familiarity had no effect on dyad performance. In the following, each of these results will be discussed in turn.

No significant difference was found when comparing the dyad to the sensitivity of better participants, therefore pairs did not significantly achieve collective benefit. This is not surprising, as Bahrami et al. (2012b) did not find that dyads achieved a collective benefit in the nonverbal condition, which the present study was modeled after. Perhaps the current exchange and quality of information is not adequate for proper integration or is insufficient information to achieve benefit. It may be of interest to explore alternative means of nonverbal communication, such as providing cumulative performance scores at the end of each block as an added means to assess a partner’s reliability.

As Bahrami et al. (2012a) and others (Wahn et al. 2017) have found, a combination of types of information exchanged may enhance collective benefit in a variety of tasks. Wahn et al. (2017) explored the effect of information type exchanged on collective benefit in a visuospatial task. They found that collective benefit could be achieved by providing either performance scores or information about a co-actor, but when provided with the combination of both information types, dyads achieved a higher performance earlier than in either condition alone. In the current paradigm, it is possible that the number of trials was an insufficient amount of time to observe collective benefit. With this in mind, the current exchange of information may not be enough to overcome individual differences and/or the limitations of nonverbal communication to achieve benefit. Perhaps providing a combination of additional information types to the nonverbal confidence rating scale would allow for dyads to achieve collective benefit within the allotted number of trials. Further experimentation would be needed in order to address either of these alternatives.

One could also potentially improve upon the confidence scale used in the current paradigm by implementing a more rigid structure. For example, each level of confidence could be assigned a more concrete classification, numerical or descriptive, therefore further aligning the dyad’s use of the scale. Bahrami et al. (2012b) found that dyads communicating verbally, as compared to nonverbally or not at all, achieved the greatest benefit and fulfilled the predictions of the WCS model. This suggests that verbal communication is most advantageous when it comes to achieving collective benefit. Under further investigation, Fusaroli et al. (2012) analyzed how dyads verbally adapted to each other and found that pairs who aligned linguistically to the task at hand locally, but more importantly, globally achieved greater benefit. As Borg (1982) proposed, the use of a scale with the addition of verbal anchors allows better for the possibility of inter-individual comparisons by creating a fixed foundation for all individuals to utilize. Therefore, adding descriptive factors to the scale may help dyads align linguistically on a more local and global level adding a fixed “verbal” concept to a nonverbal task.

Overall, participants were well calibrated, expressing a positive metacognitive sensitivity, on an individual level, which was demonstrated by the positive relationship between performance and expressed confidence level. However, dyads did not significantly accrue collective benefit to an advantage. To understand why this happens, the dynamics of the confidence rating scale were further investigated. We first looked at the difference between a pair’s confidence distributions, and found no significant relationship between confidence distribution difference and dyad sensitivity, further suggesting possible dissimilar expression of information between a dyad. In other words, knowing how often each confidence rating was used and how different this distribution was between co-actors, was not sufficient information to predict collective benefit. This is reasonable as the distribution reveals nothing about performance at a particular confidence level.

Furthermore, if a pair expresses their answers similarly, this does not imply that the information was expressed accurately or interpreted similarly. While the most extreme values along a scale leave little room for inter-individual differences in interpretation, lack of concrete descriptors allows for the possibility of diverging personal interpretations of middle values. This may make expressing one’s degree of certainty or uncertainty more difficult and the distribution of these values will only tell us how often they are used. Therefore, it is appropriate to assess the performance differences between a pair at each confidence level, which we evaluated through the CPDI measure.

The CPDI compares the alignment of a dyad based on the performance of each dyad member at each confidence level. Here, a relationship was found between CPDI and collective benefit. Thus, the more similar the usage of the confidence rating scale between a dyad, the greater the ratio of sensitivity between dyad and best member, thereby increasing the possibility of achieving collective benefit. Pescetelli et al. (2016) similarly reported a positive correlation between the average metacognitive sensitivity of individuals in a pair and dyadic performance when reporting confidence through wagers. Our results may therefore solidify the effectiveness of the rating scale as an indicative measure of performance between dyads of high sensitivity differences. This coincides with what Fusaroli et al. (2012) found in that dyads more linguistically aligned to the task at hand achieved higher performance than those who were not. However in the current study, as was observed in the Bahrami et al. (2012b) study, with the given experimental conditions no collective benefit was achieved. This suggests that the internal conceptualization of the scale between a pair is not the same, where the concept of each confidence value is not similar enough to be advantageous. More specifically, how one co-actor internalizes the value of confidence level “2” may not be similar enough to how the other co-actor internalizes that same value.

To further assess the possibility of achieving collective benefit to an advantage, we compared our data to the predictions of previously established models. Our data significantly did not fulfill the WCS model as proposed by Bahrami et al. (2010). Instead, the model overestimated dyad performance, similarly to what was found in Bahrami et al. (2012b). Despite significant deviations, the model produced a similar linear trend as our data. As the WCS model predicts dyad performance given appropriate expression of information, it would appear that information in the current study was not expressed adequately in order to achieve the model’s predictions. This may be as a result of the way in which the confidence scale was understood between dyads.

Bahrami et al. (2010, 2012a, 2012b) employed participants who were familiar with each other. This parameter was not restrictive in our study and no relationship was found between familiarity and correctness. In other words, familiar dyads were no more accurate or successful than unfamiliar dyads. This lends to a real world application of such communication measures, in that relative strangers can conceivably communicate effectively to achieve collective success. For example, such a measure could be applied across a company to achieve success among colleagues who are unfamiliar with each other or are from different departments, but need to collaborate on a task. On the other hand, Brennan and Enns (2015) found a relationship between familiarity and visibility of a partner when performing a visuospatial task. Familiar pairs were reported to outperform unfamiliar pairs when visible to each other, but when separated by a partition, familiar and unfamiliar pairs performed similarly. It is possible this effect was present in the current paradigm as dyads were separated by a partition. Further experimentation is required to investigate whether familiar dyads would have outperformed unfamiliar dyads had they been visible to each other.

In addition to evaluating factors contributing to decision-making and collective benefit, we aimed to understand the neurophysiological underpinnings of such phenomena. Investigation of neurophysiological potentials first revealed modulation of EEG signals indicating successful manipulation of contrasts. Additionally, ERPs evoked after individual response were modulated by correctness. At the joint decision, event-related potentials were affected by both correctness and conflict, where conflict over correctness produced a larger significant difference at respective electrode sites. Furthermore, an interaction between correctness and conflict indicated differential cortical representations of correctness and conflict. After presentation of feedback, joint correctness evoked significant differences in event-related potentials, which was further modified by conflict. Moreover, existing social conflict dissipated feedback-related differences evoked by correctness.

We first sought to understand the attentional processes at hand during stimulus presentation when investigating neurophysiological aspects of joint decision-making. The architecture of the attentional system could be conceptualized with control regions in fronto-parietal areas that generate modulatory signals and sensory processing areas in the occipitotemporal cortex where these modulatory signals influence ongoing visual processing (Scolari et al., 2015). Detection of the odd Gabor patch builds the fundamental information needed for this joint task through accumulation of evidence, ultimately reaching a decision, which is expressed via button press. The analysis of EEG signals at stimulus presentation revealed a significant linear relationship for contrast differences in the target stimulus at the P300 starting from 350ms after stimulus onset. P300 components are generated when the neuronal representation of the stimulus environment is changed or updated (Polich, 2007). Midline frontal-to-parietal P300 components are typical for this kind of oddball task, where these components are generated when perceptual stimulus discrimination occurs (Polich, 2007). The classical P300 P3b component is typically found after a target stimulus in oddball paradigms. It is described as having a parietal distribution on the scalp and has been linked to context updating, context closure and event categorization (Bledowski et al., 2004; Donchin and Coles, 1988; Verleger, 1988; Kok, 2001). In contrast, the predominantly frontal P3a component has been linked to an earlier stage of information gathering rather than memory and context updating (Berti, 2016). Our findings are well explained by these concepts and further validate the reliability of our experimental manipulation. Moreover, this suggests contextual working memory updates based on the presentation of different contrasts of Gabor patches.

In addition, we examined Readiness Potential and potentials following the response for the current task. Based on previous research (Hackley and Valle-Inclán, 1999; Kutas and Donchin, 1977), we expected the Readiness Potential to be modulated in this task by conditions of conflict or correctness. The cluster analysis of differences at both individual and joint responses revealed no differences in the Readiness Potential, indicating no differences in action preparation based on conflict situations or correctness of the respective interval choice.

Significant differences directly after the response indicate differential processing of conflict and correctness of responses, suggesting a social influence based on information exchange. Both conflicting situations and incorrectness elicited a late error positivity (Pe) at posterior electrode sites. A differential encoding is suggested by a significant positive cluster at frontal electrode sites for non-conflicting conditions. This is further supported by a significantly higher difference between conflicting conditions as compared to correctness conditions. This interaction indicates more pronounced differences for the error positivity, typically found at parietal electrode sites. A conflict situation does not strictly represent an error, but in contrast to the correctness of the response, one can assume that participants were aware of a mismatch. These results comply with previous findings (Orr and Carrasco, 2011; Steinhauer and Yeung, 2010), which suggest a relationship between the slow evoked Pe potential and conscious awareness of an error and further suggest Pe as a function of evidence accumulation that an error has been made. The presence of a conflict is prominent information which should be taken into account as evidence in an error-evaluation process.

Interestingly, we found differences at the ERN only for conflicting versus non-conflicting responses, strongly supporting the notion of a conflict monitoring function of this component (Botvinick et al., 2001). Nevertheless, Boldt and Yeung (2015) found only the Pe was predictive of graded changes in confidence, while the ERN failed to predict variations of confidence in single trials, suggesting an all-or-none signal. Evidence of correct-related negativity proposes the ERN component to include response elicited emotion and comparison (Vidal et al., 2000). This may implicate social context as an important factor in decision-making. The finding of an earlier deflection for response conflict, but not for correctness conditions, could be due to the fact that information that an error was made was not readily available, in contrast to the information that a conflict is present. In other studies, participants are immediately aware of an error, such as in flanker tasks where the correct response is readily available. However, in the current study the response is made after stimulus offset. Hence, we comply with theories reporting that the ERN relates to response conflict and not with theories which support it as pure error representation.

In fact, we found no error representation (correctness contrast), and this might be due to the fact that there is no clear representation of an error or feedback about an error. De Gregorio et al. (2018) suggest that representation of a correct response is necessary to elicit the ERN component. While we show presence of this ERN, a clear representation of this response may not have been formed. This is supported by the division of confidences reported, in that participants on average reported their confidence in the lowest confidence level, 53.28% of the time. Overall this suggests participants were generally uncertain in their reported decision and therefore this may be associated with an unclear representation of a correct response. Nevertheless, we see a Pe for both the correctness contrast as well as the conflict contrast, even though the information indicating the correctness of response was not available. While error-monitoring functions of the ERN and Pe are well established in literature, we provide evidence for an additional differential processing of social conflict associated with pronounced ERN and Pe.

Error-related feedback negativity and late positive potential at feedback presentation were investigated to discern differences based on individual contributions to the joint decision-making process. We could not demonstrate evidence for a modulation of the fERN by incorrect responses or conflicting individual responses. The fERN has been found in conditions where feedback is the first indication of an error (Stahl, 2010). In the current study, the presentation of feedback is not necessarily the first indication of a possible error and moreover, is not the first indication of response conflict. Once participants indicate confidences in their individual interval choice, the responses of the dyad are revealed. At this point, individuals may realize their response is in conflict with their co-actor and this may also be the first indication that one may have committed an error. Feedback is then given after the final joint decision is made.

However, a significant negative cluster, which was observed in the time window of 450 to 900 ms following feedback presentation, may represent the late positive potential (LPP). Typically, this potential is distributed over frontal-to-parietal central electrode sites (Naumann et al., 1992; Cuthbert et al., 2000; Cunningham et al. 2005; Wiens and Syrjänen, 2013), which aligns well with the location of the negative cluster found in the current study. Bradley (2009) reported that the LPP is elicited by emotional stimuli, which is interpreted to be related to enhanced attention and visual processing. Our results indicate a stronger response for conditions where the joint interval decision made by the EEG participant was incorrect, which can accordingly be interpreted as more emotionally arousing given the increased personal culpability in this decision. A significant interaction indicating higher differences for correct versus incorrect conditions when there was no conflict is particularly interesting, indicating a differential error encoding for different conflicting situations. These results can also be accounted for by interpreting conflicting situations as situations with higher perceptual uncertainty and the current design is not suitable to disentangle those two interpretations, i.e. both mechanisms could be at play.

## 5. Conclusion

The current study sought to investigate the neurophysiological correlates of joint action and collective decision-making by combining an already-established perceptual decision-making paradigm with EEG. Similar to Bahrami et al. (2012b), we found that dyads were overall unable to achieve collective benefit to an advantage. While neither the WCS nor the optimal models provided a significant fit for observed behavior, the WCS model predicted the same dependence on individual performance similarity with higher performance offset. Through investigating the relationship between confidence performance difference index (CPDI) and collective benefit, we sought to better understand the confidence rating scale as a means of communication. Our results indeed suggest that the CPDI could be a predictive measure of collective benefit.

In addition, our results for the neurophysiological data suggest the presence of attentional, error-evaluation or social conflict-monitoring processes in each investigated phase in the current paradigm. In particular, in the stimulus presentation phase, a contextual working memory update is possibly represented by the linear relationship of contrast differences and the P300. When a joint interval decision is indicated by a button press, a social conflict due to divergent individual decisions may be represented by a frontal error-related response negativity in the response phase. Subsequently, incorrect decisions and social conflict may be represented in a late error positivity potential. Similarly in the feedback phase, a late positive potential elicited after the presentation of feedback by incorrect and conflicting situations is indicative of emotional arousal and an allocation of attentional resources.

In summary, our results suggest that response- and feedback-related components and potentials elicited by an error-monitoring system may differentially integrate socially conflicting information exchanged during the joint decision-making process. The current study serves as an initial exploratory step towards understanding the underlying mechanisms involved in dyadic interaction and exchange of information. Further research could facilitate understanding of the suggested underlying mechanisms and solidify effective communication techniques for dyads of diverging sensitivities.

## 6. Conflict of Interest

The authors declare that the research was conducted in the absence of any commercial or financial relationships that could be construed as potential conflicts of interest.

## 7. Author Contributions

Study Design: PK, BW, KGB, PB, A-NS, and AS. Data Acquisition: KGB, PB, A-NS, and AS. Experimental Task Programming: KGB, PB, and A-NS. Behavioral Data Analysis: KGB, PB, and A-NS. EEG Data Analysis: AS. Literature Research: KGB, PB, A-NS, and AS. Manuscript Writing: KGB, PB, A-NS, and AS. Manuscript Revisions: KGB, PB, A-NS, AS, PK, and BW.

## 8. Funding

We acknowledge the support of a postdoc fellowship of the German Academic Exchange Service (DAAD) awarded to BW. Furthermore, we acknowledge the support by H2020 –H2020 FETPROACT-2014 641321 – socSMCs for PK and the support from the Deutsche Forschungsgemeinschaft (DFG), and the Open Access Publishing Fund of Osnabrück University.

## 9. Acknowledgements

We would like to thank Paola Suarez-Ramirez and Raul Sulaimanov for their contributions to data collection.

