## Supplementary Material for "Neurophysiological correlates of collective perceptual decision-making"

### Supplementary Figures

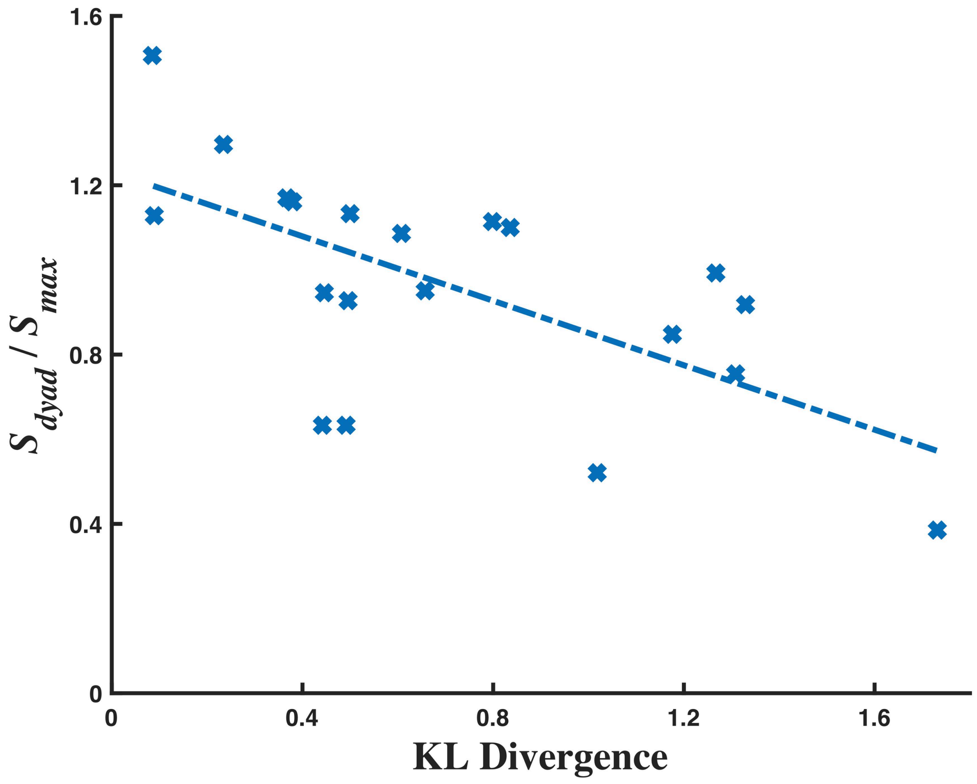


**Supplementary Figure 1.** The relationship between KL Divergence and collective benefit ($S_{dyad}/S_{max}$) per dyad (regression fit, blue dashed line).

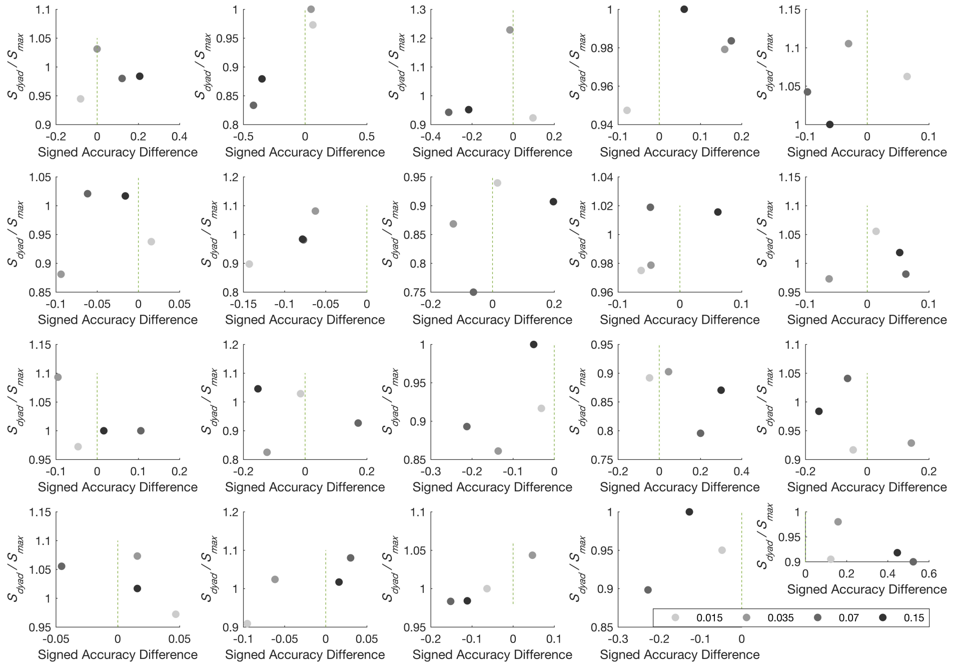


**Supplementary Figure 2.** The relationship between signed accuracy difference and collective benefit for each difficulty level on a per dyad basis. Gradient data points from light to dark signify the most difficult level to the easiest level. A signed accuracy difference of 0 indicates identical accuracy for both dyad members. A value greater than one for the $S_{dyad}/S_{max}$ ratio signifies a collective benefit to an advantage.
